# Single-cell transcriptomic atlas of mouse oocyte development from growth to ovulation

**DOI:** 10.64898/2026.03.11.710939

**Authors:** Sining Zhou, Zhenshe Lin, Yiming Liu, Chang Liu, Xinyu Yang, Meichi Yue, Hong Wu, Rui Liu, Huinan Xu, Xiaoxi Zhou, Qian Liu, Zhe Zhang, Liyun Xiao, Jingxuan Zou, Jiamei Liu, Yuan Li, Shijie Hao, Yijun Ruan, Xun Xu, Chuanyu Liu

## Abstract

Oocyte growth and maturation depend on tightly coordinated programs within oocytes and their surrounding granulosa cells, yet defining the transcriptional continuum of growing oocytes has been challenging due to their large size and the limitations of current droplet-based and spatial transcriptomic platforms. Here, we optimized a high-density array–based platform called Stereo-cell for high-throughput dual-modality profiling of large mouse oocytes, allowing for both the preservation of morphology and transcript capture. By integrating unsupervised transcriptomic clustering with cell morphological features, we delineated successive temporal windows from growing oocytes to metaphase II and uncovered stage-linked shifts from early programs toward later programs. We also profiled the ovarian somatic fraction, reconstructed granulosa-cell subtype relationships, and placed ovarian cell states in tissue context using single-cell–resolution spatial data.

## Introduction

Folliculogenesis is a coordinated developmental process in which endocrine cues from the hypothalamic–pituitary–gonadal axis are interpreted by ovarian somatic cells and translated into local paracrine and contact-dependent signals that sustain oocyte growth and competence acquisition^1^. Over the past decade, single-cell and spatial transcriptomic technologies have greatly expanded our understanding of ovarian cell states and niche organization^2^, enabling the creation of atlases of cycling and aging mouse ovaries, primordial follicle assembly, adult human ovarian cell landscapes, and spatiotemporal dynamics during ovulation^3–8^. Yet, a central gap remains along the oocyte “growing” continuum. Growing oocytes undergo dramatic size increases and continuous transcriptional remodeling. However, their large dimensions hinder efficient capture by droplet-based platforms, and ovarian spatial transcriptomic datasets often lack sufficient effective resolution to disentangle oocyte signals from surrounding granulosa cells (GCs)^5,6^. As a result, profiling of growing oocytes has often relied on low-to-moderate throughput, tube-based protocols, with staging and grouping typically determined by manual morphological assessment—an approach that introduces subjectivity and limits continuous, low-batch sampling across developmental stages^9–13^.

Here, we address these limitations using Stereo-cell, a high-density array–based platform for spatially resolved single-cell transcriptomics with integrated imaging. We applied and optimized the platform for large oocytes to obtain high-throughput, morphology-linked oocyte profiling^14^. By refining oocyte pretreatment to improve morphological preservation while maintaining transcript capture, we enabled joint interpretation of transcriptional state and cell morphological features, which supports assigning oocytes into successive temporal windows along growth and maturation with minimal manual bias. In parallel, we profiled the ovarian somatic fraction and single-cell–resolution spatial data to place granulosa subtypes into tissue context and to examine stage-resolved oocyte–granulosa communication programs. Together, this work provides an integrated resource for dissecting oocyte-intrinsic programs and microenvironmental interactions during the previously under-characterized transition from growth to maturation.

## Results

### Study design for single-cell RNA-seq across mouse oocyte growth

To capture the transcriptomic changes of mouse oocytes from primary to preovulatory follicles, we collected ovaries from postnatal day (P)14, P21, and P49 C57BL/6 mice, as well as from hormone-primed (HP; PMSG + hCG) P21 oviducts. After tissue dissociation, oocytes were manually picked, and the remaining cell suspension was passed through a filter to obtain ovarian somatic cells. Oocyte transcriptomes were profiled using the Stereo-cell platform, whereas ovarian somatic cells were profiled using the DNBelab C4 droplet-based single-cell RNA-seq system (**Fig. 1a**).

**Fig. 1.**
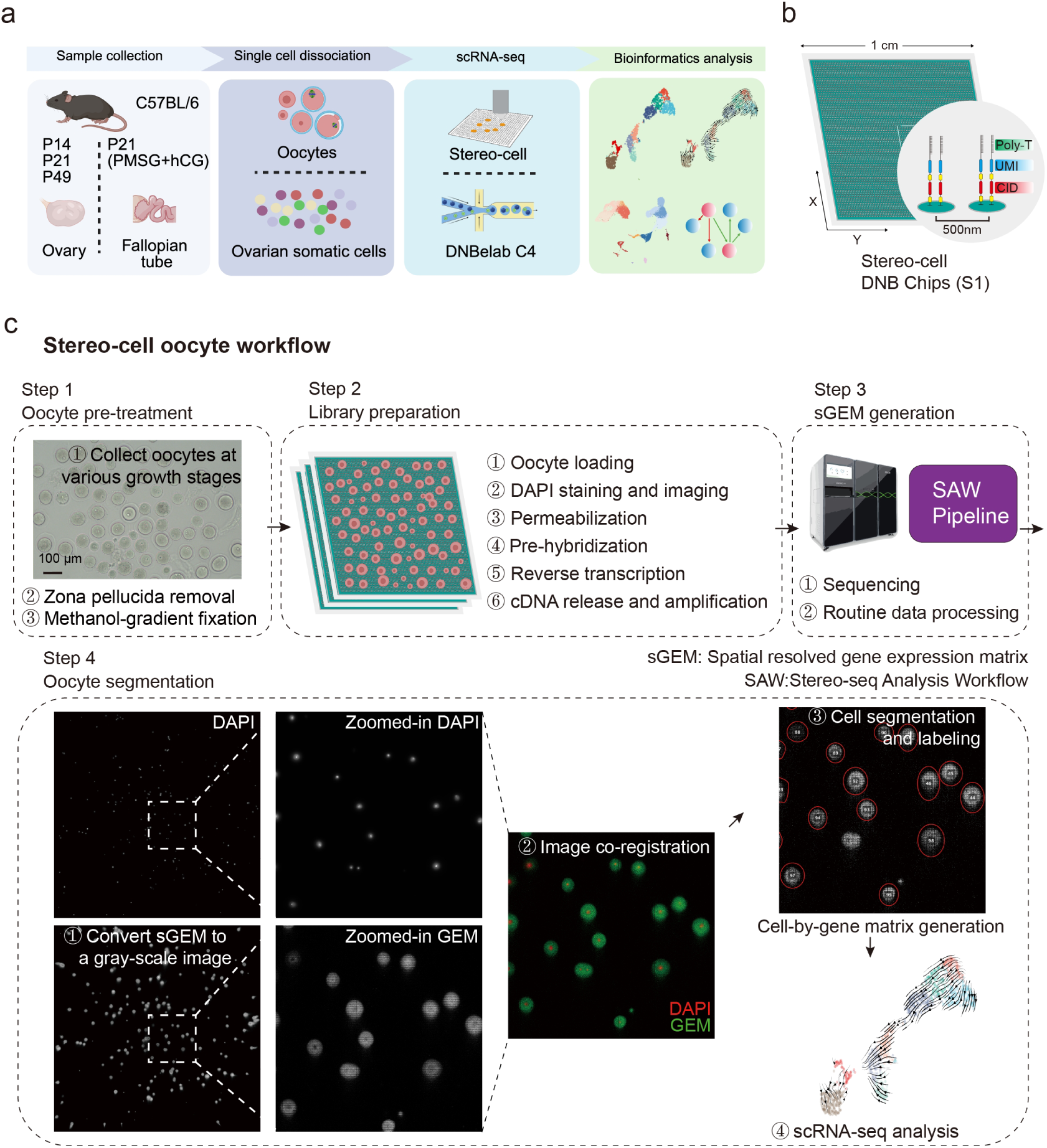
Study design and Stereo-cell workflow for dual-modality profiling of mouse oocytes. **a**, Overview of this work. Oocytes were collected from C57BL/6 mice at P14, P21, and P49, with an additional cohort obtained from hormonally primed P21 mice (PMSG + hCG). Oocytes were manually isolated for Stereo-cell profiling, while the ovarian somatic-cell fraction was processed in parallel using the DNBelab C4 droplet-based scRNA-seq platform. **b**, Schematic of the Stereo-cell patterned DNA nanoball (DNB) chip (S1). DNBs are arranged on the chip surface and functionalized with capture probes containing poly(dT), unique molecular identifiers (UMIs), and spatial coordinate barcodes (CIDs). **c**, Stereo-cell oocyte workflow. Step 1, oocyte pre-treatment including collection of denuded oocytes at different growth stages, zona pellucida removal, and methanol-gradient fixation (scale bar, 100 μm). Step 2, on-chip library preparation including oocyte loading, DAPI staining and imaging, permeabilization, pre-hybridization, reverse transcription, and cDNA release and amplification. Step 3, sequencing and routine data processing to generate spatially resolved gene expression matrices (sGEM) using the SAW pipeline. Step 4, oocyte segmentation and image–transcriptome integration: sGEM is converted to a gray-scale image and co-registered with on-chip DAPI images, followed by cell segmentation and labeling to generate a cell-by-gene expression matrix for downstream single-cell transcriptomic analysis. Panel a was created in BioRender.

The Stereo-cell multi-omics technology is built on patterned DNA nanoball (DNB) chips that follow the Stereo-seq design (**Fig. 1b**)^14,15^. Each chip contains DNBs arranged at 500 nm intervals and functionalized with capture probes carrying poly(dT) tails, unique molecular identifiers (UMIs), and spatial barcodes (coordinate IDs [CIDs]), enabling high-throughput, spatially resolved transcript capture. Because Stereo-cell is not limited by cell size, it is well suited for high-throughput profiling of large cells such as oocytes. In addition, the platform supports on-chip staining, allowing simultaneous acquisition of transcriptomic and morphological information. Thus, for each oocyte we can obtain its transcriptome together with cell size and DNA staining patterns, which facilitates more accurate assignment of developmental stage.

In this study, we further optimized the pre-treatment steps for applying Stereo-cell to mouse oocytes. We collected and isolated morphologically intact denuded oocytes and removed the zona pellucida to enhance permeabilization according to an established protocol^16^. A methanol-gradient fixation strategy was applied to reduce the deformation that can be caused by rapid dehydration in high concentrations of methanol. After these pre-treatment steps, oocytes were loaded onto Stereo-cell chips. Subsequent library construction, sequencing, and data processing followed our previously published Stereo-cell workflow. On-chip DAPI imaging was co-registered with the post-sequencing spatially resolved gene expression matrix (sGEM), providing a unique index for each oocyte and linking its morphology to its transcriptome. This image–transcriptome integration enables high-throughput, dual-modality (morphology plus transcriptome) profiling of mouse oocytes (**Fig. 1c**).

### Integrated transcriptomic and morphological staging of mouse oocyte maturation

Across 10 Stereo-cell libraries, we obtained transcriptomes for 1,051 oocytes. Unsupervised clustering of the integrated dataset identified nine clusters (C0–C8) (**Fig. 2a and Supplementary Fig. 1a**).

**Fig. 2.**
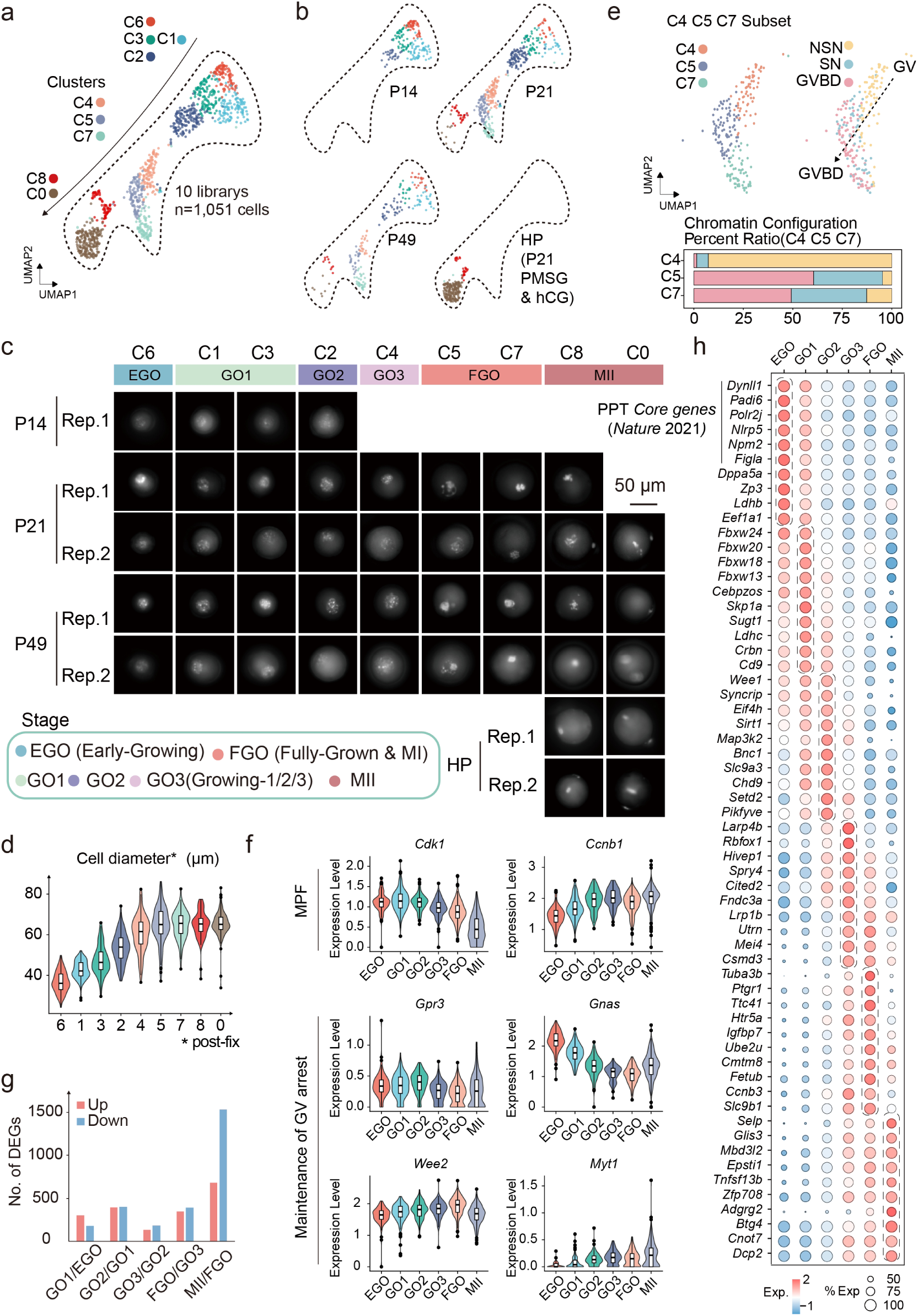
Integrated transcriptomic and morphological staging of mouse oocyte maturation. **a**, UMAP of 1,051 oocytes integrated from 10 Stereo-cell libraries, colored by unsupervised clusters (C0–C8). **b**, UMAP shown separately for oocytes collected from P14, P21, and P49 ovaries and from HP oviducts. **c**, Representative on-chip DAPI images of oocyte nuclei across clusters and sampling conditions (P14, P21, P49, HP; replicates indicated). Clusters are grouped into developmental stages (EGO, GO1, GO2, GO3, FGO, and MII) as indicated. Scale bar, 50 μm. **d**, Violin plots of oocyte diameter across clusters. **e**, Subset analysis of clusters C4, C5, and C7. Left, UMAP colored by clusters; right, UMAP colored by chromatin configuration states (NSN, SN, and GVBD). Bottom, percentage of chromatin configurations in each cluster. **f**, Violin plots showing relative expression of *Cdk1*, *Ccnb1*, *Gpr3*, *Gnas*, *Wee2*, and *Myt1* across clusters. **g**, Numbers of upregulated and downregulated differentially expressed genes (DEGs) between consecutive stages. **h**, Dot plot of representative stage-signature genes across the annotated stages (EGO, GO1, GO2, GO3, FGO, and MII). Dot color indicates scaled expression and dot size indicates the fraction of cells expressing the gene.

Mice at postnatal day 14 are commonly used to isolate immature follicles^16^. In our work, P14 oocytes were predominantly located in clusters C6, C3, and C1, with a small fraction in C2. In contrast, oocytes collected from the HP oviducts were almost exclusively assigned to clusters C8 and C0. These observations suggest that clusters C6, C3, and C1 represent early-stage oocytes, whereas clusters C8 and C0 correspond to late-stage oocytes. Oocytes from P21 and P49 ovaries were distributed across all clusters, consistent with previous reports that ovaries at these time points contain oocytes spanning the full range of growth stages (**Fig. 2b**)^17^.

By labeling nuclear DNA with DAPI, we captured stage-dependent changes in nuclear configuration and cell size during oocyte development and used these features to define distinct developmental stages (**Fig. 2c–d**). Oocytes in clusters C8 and C0, derived from both unstimulated ovaries and HP oviducts, exhibited first polar body extrusion or separation of homologous chromosomes, consistent with metaphase II (MII) arrest or late meiotic maturation along the metaphase I (MI)-to-MII continuum. Cells in C5 and C7 displayed partially or predominantly condensed (pSN/SN) or germinal vesicle breakdown (GVBD)-like nuclei and transcriptomes close to full maturation (**Fig. 2e**), so we classified them as fully grown oocytes (FGO)^18^. In contrast, C6 contained oocytes with the smallest diameters, indicating the least advanced developmental stage; we therefore defined this cluster as early-growing oocytes (EGO). Oocytes in C1 and C3, which showed highly correlated expression and had larger diameters but similar nuclear morphology to C6 (**Supplementary Fig. 1b**), were annotated as growing oocyte-1 (GO1). Comparison with published data on oocyte diameter measurements further suggested that EGO oocytes mainly derive from late primary follicles, whereas GO1 oocytes correspond to early secondary follicles (**Supplementary Fig. 1c-e**)^19^. C2 showed a further increase in size relative to GO1 and was more strongly correlated with C4/C5/C7 at the transcriptomic level; this cluster was therefore defined as GO2. Cluster C4 lay near the more mature clusters (C5/C7) in UMAP space but retained a diffuse non-surrounded nucleolus (NSN) chromatin pattern. We designated this cluster GO3, as it represents a candidate transition state from growing to fully grown oocytes. Cell diameter increased from C2 through C5, then plateaued in C7, C8, and C0 (**Fig. 2d, e**).

After defining developmental stages by integrated transcriptomic and morphological features, we inspected canonical regulators of prophase I arrest and meiotic competence. The maturation promoting factor (MPF) core components *Cdk1* and *Ccnb1* were broadly expressed across growing stages, with *Ccnb1* gradually accumulating with maturation, whereas the cAMP–GV arrest axis (*Gpr3* and *Gnas*) was already readily detectable in early stages. Consistent with increasing meiotic competence, the inhibitory kinases *Wee2* and *Myt1* increased toward later stages, indicating strengthened inhibitory control over *Cdk1* activity as oocytes approach full growth, consistent with maintaining prophase I arrest in the face of progressive MPF accumulation (**Fig. 2f**)^20,21^. In parallel, transcriptional activity progressively declines during late oocyte growth and is essential for oocyte chromatin reorganization^22^. Accordingly, the CTD kinase *Cdk9*, which phosphorylates RNA polymerase II to promote elongation, was higher in C6–C4 (EGO–GO3) but declined from C5 onward (FGO to MII), whereas the opposing CTD phosphatase *Ctdp1* remained comparatively stable across stages (**Supplementary Fig. 1f**). These observations suggest that transcriptional elongation may be attenuated between GO3 and FGO^23,24^.

Together, these analyses show that unsupervised clustering of Stereo-cell transcriptomes, combined with on-chip nuclear morphology and cell size, enables robust annotation of oocyte developmental stages.

### Transcriptional signatures across oocyte maturation

Using Seurat’s FindAllMarkers, we identified stage-specific highly expressed genes and quantified upregulated and downregulated genes between consecutive stages (**Fig. 2f**). From the highest log2 fold-change rankings, we selected 10 representative genes per stage (**Fig. 2g**).

EGO highly expressed core transcription factors and markers of the primordial-to-primary follicle transition (PPT), including *Dynll1*, *Padi6*, *Polr2j*, *Nlrp5*, *Npm2*, *Figla* and *Zp3*, further aligning EGO with primary follicle oocytes^19^. GO1 preserved some PPT genes and showed peak expression of Fbxw family members (*Fbxw24/20/18/13*) rising from EGO, highlighting a stage-specific enrichment of SCF/F-box–associated ubiquitin pathway components^25^. GO2 retained GO1-enriched genes, while *Syncrip* (a post-transcriptional regulator)^26^, *Eif4h* (a eukaryotic translation initiator)^27^, *Map3k2* (MAPK upstream kinase)^28^ and *Chd9* (an enhancer of chromatin accessibility)^29^ reached peak expression at this stage.

Interestingly, GO3 versus GO2 showed the fewest differentially expressed genes (DEGs) among adjacent stages, yet many expression patterns clearly split at this boundary. Many genes that were progressively induced across GO1–GO2 reversed course and dropped at GO3, whereas genes enriched in GO3/FGO/MII began to rise rapidly at GO3. This pattern is consistent with unsupervised clustering, in which EGO–GO2 group together and GO3–FGO cluster together, and points to a major shift in the oocyte transcriptional program at the GO2–GO3 switch. GO3-enriched genes included *Larp4d* (mRNA stability) and *Rbfox1* (alternative splicing)^30,31^, which were already rising in GO2, and GO3-onset genes such as *Utrn* (involved in cytoskeletal organization) and *Mei4*^32,33^ (involved in meiotic recombination), indicating active mRNA storage, splicing regulation, and cytoskeletal dynamics, which positions GO3 as a transitional window between growing and fully grown.

FGO is a mature pre-resumption stage characterized by the NSN-to-surrounded nucleolus (SN) transition and global transcriptional silencing. Its enriched genes rose from GO3, including *Ptgr1*, an NADPH-dependent prostaglandin reductase that inactivates arachidonic acid (ARA)-derived eicosanoids and lipid peroxides^34^. Prior proteomics observed a decline of ARA from germinal vesicle (GV) to GVBD^35^, and in our dataset *Ptgr1* peaked in FGO, suggesting that *Ptgr1* abundance may support terminal ARA metabolism and facilitate ARA clearance during late meiotic maturation.

In MII, we observed the largest number of downregulated genes, consistent with extensive maternal mRNA degradation. However, many genes showed relatively high transcript levels, as also noted in previous studies^36^. Given transcriptional quiescence after FGO (particularly after GVBD), these transcripts are unlikely to be newly synthesized but instead maintained for storage or translation to execute essential functions (**Fig. 2f**). This is likely the case for degradation regulators *Btg4*, *Cnot7*, and *Dcp2*, which are critical for maternal mRNA clearance and oocyte developmental competence^37,38^.

To independently validate these stage-specific signatures, we performed single-sample gene set enrichment analysis (ssGSEA) on a published bulk RNA-seq dataset (GSE70116) of oocytes at different time points of follicular growth^39^. In this dataset, scores derived from our EGO-enriched genes progressively decreased from NGO stage (non-growing oocytes, 10–40 µm) to FGO stage (fully-growing oocytes, >70 µm), whereas scores for the FGO signature increased with maturation, indicating that our Stereo-cell signatures faithfully capture the transition from immature to mature oocytes. GO1/GO2/GO3 signature scores also rose to varying degrees after NGO, broadly mirroring our single-cell trends. Bulk profiling, however, lacked the resolution to distinguish the fine-grained differences we observed between GO1, GO2, and GO3 (**Supplementary Fig. 1g**).

Together, these results outline distinct transcriptional signatures for each stage: EGO is dominated by PPT-associated, GO1–GO2 by progressive growth with enhanced protein homeostasis and signaling; FGO by NSN-to-SN transition and strengthened post-transcriptional control; and MII by reinforced maternal mRNA clearance.

### Co-expression gene modules and regulatory programs during oocyte maturation

To dissect transcriptomic changes during oocyte growth and infer developmental trajectories, we applied two root-independent approaches, scTour and CytoTrace2^40,41^. The scTour-derived trajectory closely matched the Seurat clustering, supporting the inferred developmental direction. CytoTrace2 scores placed all oocytes within a high developmental-potential range (score > 0.83) and gradually increased from immature to meiotically mature states, forming a continuous early-to-late gradient (**Fig. 3a, b**).

**Fig. 3.**
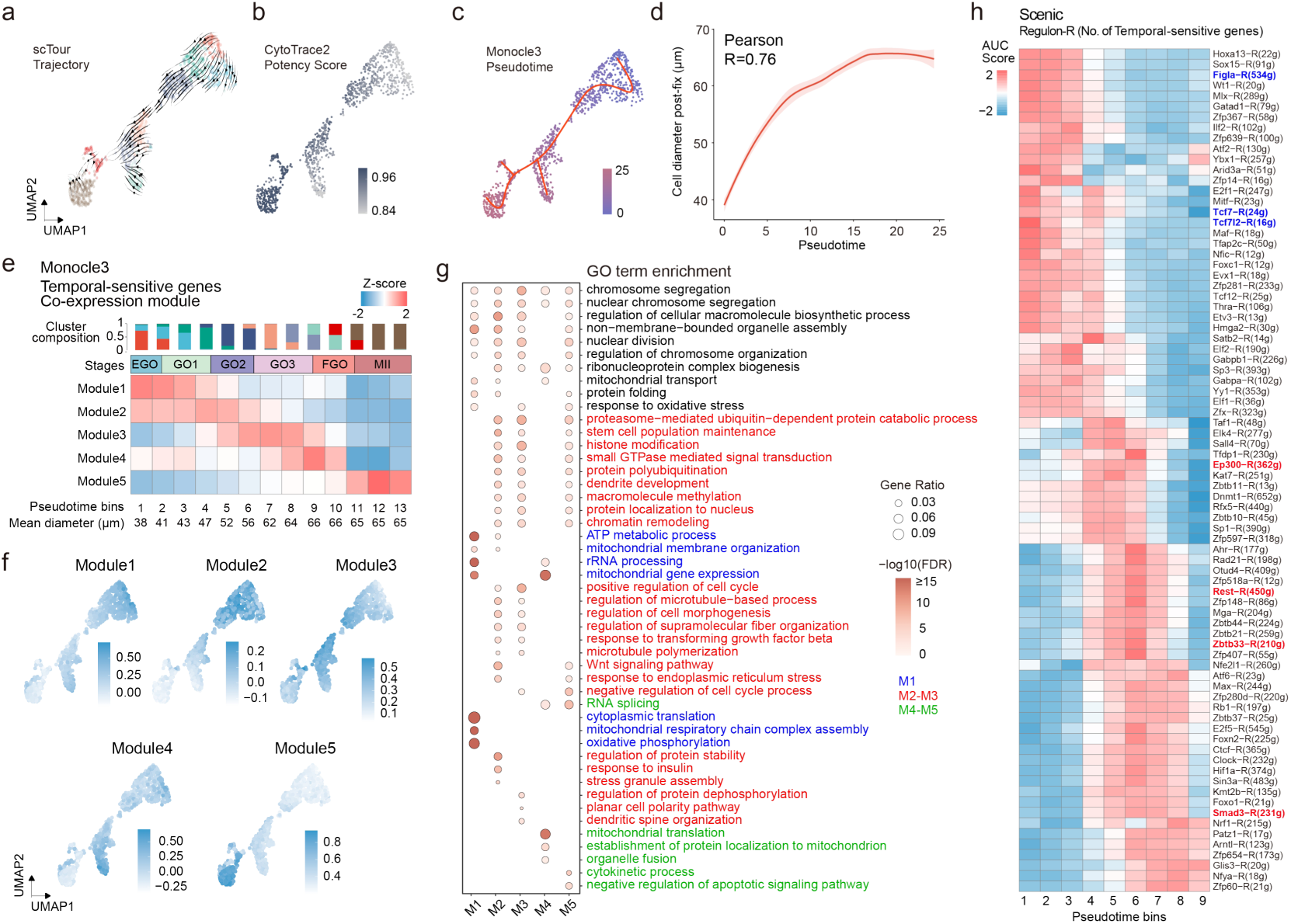
Co-expression gene modules and regulatory programs during oocyte maturation. **a**, scTour-inferred trajectory projected onto the UMAP embedding of oocytes; arrows indicate the inferred developmental direction. **b**, CytoTrace2 developmental potency score overlaid on the UMAP. **c**, Monocle3 pseudotime ordering visualized on the UMAP with the learned principal graph. **d**, Relationship between Monocle3 pseudotime and oocyte diameter, with Pearson correlation shown. **e**, Heatmap of pseudotime-sensitive genes (Moran’s I >0.2) grouped into five co-expression modules (M1–M5). Cells are evenly binned into 13 pseudotime bins; values are z-scored across bins. **f**, UMAP feature plots showing module scores for M1–M5. **g**, GO term enrichment analysis for genes in modules M1–M5. **h**, SCENIC regulon activity (AUC) across pseudotime bins (1–9). Numbers in parentheses indicate regulon size.

We next performed Monocle3 pseudotime analysis^42^, setting C6 as the root. Cell diameter increased rapidly and then plateaued along pseudotime, consistent with morphology-based staging (**Fig. 3c, d**). Using Moran’s I > 0.2, we identified 3,211 pseudotime-sensitive genes and grouped them into five co-expression modules (M1–M5) with findgenemodules. After evenly dividing cells into 13 pseudotime bins, module heatmaps revealed continuous yet stage-biased expression across neighboring bins. Modules M1–M4 corresponded to genes upregulated across EGO→GO1, GO1→GO2, GO2→GO3, and GO3→FGO transitions, respectively, whereas M5 contained transcripts that remained abundant at MII (**Fig. 3e, f**).

GO analysis highlighted distinct biological programs (Fig. 3g). M1, predominant in EGO, was enriched for mitochondrial respiration/oxidative phosphorylation. Consistent with this, most nuclear-encoded mitochondrial genes showed generally higher transcript abundance in EGO (primary follicle oocytes). This early peak was also observed in an independent dataset (CRA001613)^12^, where primary-follicle oocytes (T3a–T3b) exhibited similarly elevated expression (**Supplementary Fig. 2a**)^36^. This pattern aligns with the strong enrichment of oxidative phosphorylation terms in M1, suggesting that the primary-follicle stage represents an early window of metabolic activity demand. In contrast, mitochondrial genome–encoded transcripts displayed an opposite trend, with relatively higher levels at MII (**Supplementary Fig. 2b**).

M2 peaked at late GO1 (bins 4–5), and M3 peaked around the GO2–GO3 transition (bins 6–7), marking two major windows of coordinated gene regulation. Relative to M1, M2/M3 reflected a functional shift toward protein homeostasis and stress responses, signaling and cell-cycle/stemness control, epigenetic/chromatin regulation, and cytoskeleton/morphogenesis-related processes. Within this framework, M2 was more associated with protein stability and growth factor/stress-related responses, whereas M3 was enriched for structural and polarity remodeling, consistent with progressive reorganization during GO2–GO3.

Modules active around GO3–FGO and beyond were biased toward post-transcriptional and mitochondrial events. M4, rising from GO3 and peaking in FGO, was enriched for RNA splicing and mitochondrial translation/import, consistent with intensified post-transcriptional control of maternal mRNAs around meiotic resumption. M5 transcripts remained abundant at MII and were enriched for RNA processing, cytokinesis-related processes and anti-apoptotic programs, suggesting retained RNAs support late meiotic dynamics and oocyte viability.

SCENIC inference of regulon activity restricted to bins 1–9 (excluding bins 10–13, which correspond to meiotic resumption with minimal nascent transcription) recapitulated the module-defined maturation trajectory and highlighted two major transitions^43^. Around bins 3–4, EGO/GO1-associated early oocyte growth programs rapidly attenuated (e.g., *Figla* sharply declining from bin 4 onward), whereas the GO2–GO3 interval coincided with a broad rise of repression-associated regulons (e.g., *Rest* and zinc-finger/BTB programs) ^44^. WNT and TGFβ signaling are established pathways in oocytes^12^, with WNT reported to become activated around the transition into secondary follicle growth^45^. In our data, WNT-linked regulons (*Tcf7*/*Tcf7L2*) peaked earlier than *Smad3*^46^, suggesting a temporal ordering of these developmental inputs along oocyte growth.

Moreover, we also notice that several GO terms spanned multiple modules. Histone modification, for example, was significantly in M2, M3, and M5 (**Supplementary Fig. 2c**). We therefore extracted histone-modification genes from the pseudotime-sensitive set and grouped them by the stage of maximal expression, generating six histone-modification modules whose scores tracked their assigned stages (**Supplementary Fig. 2d**). At the transcript level, EGO–GO2 stages showed elevated expression of regulators associated with an active chromatin state, including *Setd1b* (H3K4 methyltransferase) ^47^, *Kdm1b* (H3K4me1/2 demethylase)^48^, *Setd2* (H3K36me3 methyltransferase)^49^, and the acetyltransferase co-activators *Ep300* and *Crebbp*^50^. After GO2, this profile shifted, with a progressive decrease of transcripts linked to chromatin opening and transcriptional support, alongside increased expression of factors associated with chromatin compaction and repression, including *Setdb1* (H3K9 methyltransferase)^51^, the *Hdac2*/*Sin3a* deacetylase complex^52^, and multiple PRC1/PRC2 components (*Pcgf6*, *Bmi1*, *Rnf2*, *Suz12*, *Eed*, *Ezh2*, *Phf1*)^53,54^, many of which remained high at the transcript level in MII (**Supplementary Fig. 2e**). Consistently, inferred stage-restricted activity patterns of chromatin-regulatory networks paralleled this gene-level transition. Consistently, SCENIC-inferred regulon dynamics for chromatin-associated regulators (e.g., Ep300-R and Sin3a-R) were stage-restricted and paralleled this transcript level shift.

Building on the pseudotime framework, these analyses delineate stage-specific programs through co-expression modules and regulatory dynamics and further pinpoint GO2–GO3 as a key transitional window. Within this transition, coordinated programs shift toward strengthened repressive chromatin activities, consistent with the establishment of transcriptional silencing as oocytes approach full growth.

### Molecular and cellular signatures of GC subtypes

While the oocyte trajectory delineates stage-specific biological programs active during oocyte growth, it does not capture the full in-follicle context in which somatic cells continuously shape oocyte maturation. To place oocyte development in this follicular microenvironment, we profiled the ovarian somatic-cell fraction in parallel using DNBelab C4 scRNA-seq. We annotated 47,412 cells that passed standard quality control using marker genes reported in previous ovarian single-cell atlases, identifying five major cell types comprising 16 subtypes (**Supplementary Fig. 3a, b**)^4–6^. Among these, GCs—the principal somatic component of ovarian follicles—communicate intimately with oocytes via paracrine and contact-dependent interactions and are essential for follicle growth and oocyte maturation. To refine GC heterogeneity for subsequent oocyte–GC signaling analyses, we regressed out cell-cycle effects and further resolved GCs into seven subtypes based on subtype-enriched markers (**Fig. 4a and Supplementary Fig. 3c**), establishing a subtype-resolved GC framework for downstream communication analyses.

**Fig. 4.**
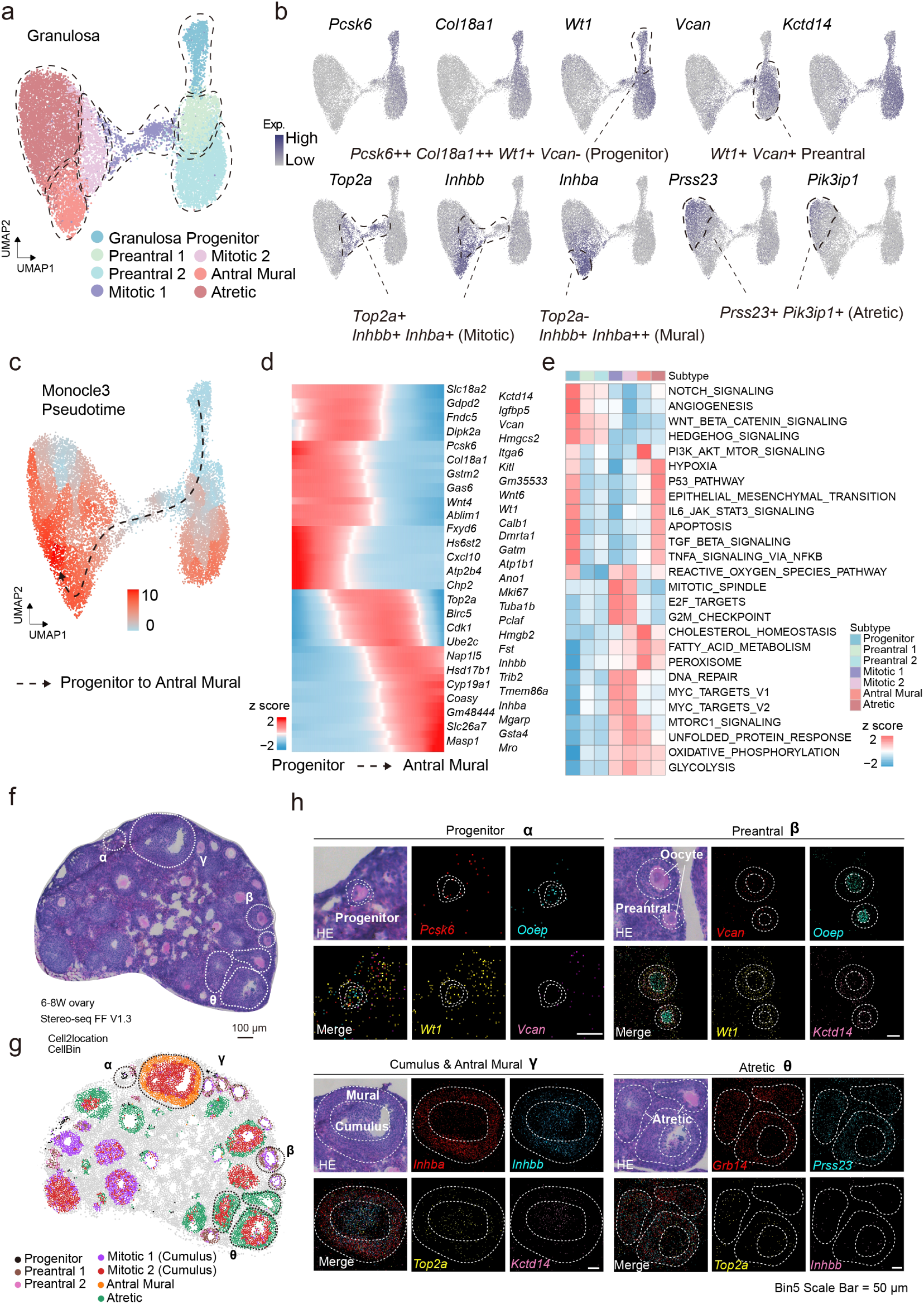
GC subtypes and spatial mapping across follicular development. **a**, UMAP of GCs, colored by seven GC subtypes (Progenitor, Preantral 1, Preantral 2, Mitotic 1, Mitotic 2, Antral Mural and Atretic). **b**, Feature plots of representative subtype markers on the UMAP. **c**, Monocle3 pseudotime trajectory inferred for GCs, with the principal graph overlaid and direction indicated from progenitor toward antral mural cells. **d**, Heatmap of representative genes showing coordinated expression changes along the progenitor-to-mural trajectory (expression shown as z-scores). **e**, Heatmap of Hallmark ssGSEA scores across granulosa subtypes (z-scored per gene set). **f**, H&E image of a Stereo-seq FF V1.3 mouse ovary section (6-8 weeks old), with representative regions (α–θ) indicated. Scale bar, 100 μm. **g**, Cell2location-based spatial mapping of GC subtypes at cell-bin resolution. **h**, Zoom-in views of representative regions (α, β, γ, and θ) showing H&E morphology and spatial expression of selected marker genes. Scale bar, 50 μm.

Within the preantral stage, we identified three GC subpopulations: Progenitor, Preantral 1, and Preantral 2. Progenitor showed higher expression of early markers *Pcsk6*, *Col18a1*, and *Gatm* and displayed a *Wt1*^+^ *Vcan*^−^ profile, consistent with pGCs (progenitor granulosa cells) described in human ovarian atlases. In contrast, Preantral 1 and Preantral 2 were characterized by *Wt1*^+^ *Vcan*^+^ expression, consistent with progression along the pGC differentiation trajectory^7^. Although these two subsets were transcriptionally similar, Preantral 2 expressed higher levels of *Igfbp5*^55^, suggesting increased engagement of IGF-related processes, including proliferation, differentiation, and apoptosis. Preantral 1 was positioned adjacent to the mitotic compartment, indicating that it may represent an intermediate state during the transition from preantral to proliferative GCs (**Fig. 4b and Supplementary Fig. 3c**).

Mitotic GCs were further separated into Mitotic 1 and Mitotic 2. Both subsets expressed proliferation-associated genes such as *Top2a* and *Hmgb2*, whereas Mitotic 1 retained higher expression of preantral features (e.g., *Kctd14* and *Vcan*), suggesting greater similarity to the preantral state. By contrast, Mitotic 2 showed reduced expression of these preantral genes and increased expression of mural-associated genes, including *Nap1l5*, *Inhbb*, and *Inhba*, indicating a shift toward a more differentiated direction. Cell-cycle scoring showed elevated S-phase scores in both mitotic subsets, while Mitotic 1 exhibited higher G2/M scores, consistent with a higher fraction of cells in late G2/M progression (Supplementary Fig. 3d). Relative to mitotic GCs, Antral Mural GCs were marked by *Mro* and showed higher *Inhba* expression (Supplementary Fig. 3c), whereas Atretic was marked by elevated *Cfh*, *Prss23*, and *Grb14* and relatively specific expression of *Pik3ip1*, consistent with an atresia-associated program (**Fig. 4a, b**).

Collectively, these subtypes delineate a continuous transcriptional progression of GC development, from *Pcsk6*^++^ *Col18a1*^++^ *Wt1*^+^ *Vcan*^−^ progenitor to *Wt1*^+^ *Vcan*^+^ preantral states, followed by a *Top2a*^+^ *Inhbb*^+^ *Inhba*^+^ proliferative/differentiation transition. This progression ultimately culminated in divergence toward *Top2a*^−^*Inhbb*^+^ *Inhba*^++^ mural GCs or an atretic branch (**Fig. 4b**). To further validate this inferred progression, we performed Monocle3 pseudotime analysis on GCs and reconstructed a continuous trajectory from the progenitor compartment through mitotic intermediates toward the antral mural state (**Fig. 4c**). Differentially expressed genes along pseudotime recapitulated the subtype transitions defined above, with early progenitor/preantral markers gradually decreasing and mural-associated genes progressively increasing. In addition to canonical markers, this analysis highlighted additional genes with coordinated temporal dynamics, including *Fndc5*/*Gdpd2* (following *Vcan*-like trends) and *Mgarp*/*Masp1* (increasing in parallel with *Inhba*) (**Fig. 4d**).

Hallmark ssGSEA analysis further revealed clear functional stratification among these subtypes (**Fig. 4e**). Progenitors were enriched for developmental and niche-communication programs (e.g., NOTCH/WNT/HEDGEHOG and angiogenesis)^56,57^, with detectable inflammatory signaling (TNFA–NFκB/IL6–STAT3), whereas preantral cells showed intermediate activities. Antral mural cells were dominated by metabolic maturation, including lipid/cholesterol and mitochondrial programs (fatty-acid metabolism, cholesterol homeostasis, and OXPHOS) with enhanced PI3K–AKT–mTOR signaling. Mitotic cells showed a sharp shift from progenitor/preantral states toward proliferation (E2F/G2M, mitotic spindle, and DNA repair) while concomitantly increasing mural-like metabolic/proteostasis programs (lipid/cholesterol metabolism, mTORC1/OXPHOS, and UPR), consistent with a proliferative-to-mural transition along the inferred trajectory. Atretic cells were enriched for stress responses (P53, hypoxia/ROS, and UPR) with elevated apoptosis and EMT (**Fig. 4e**). Although atretic and progenitor cells shared enrichment in some Hallmark categories, their gene-level signatures diverged: atretic APOPTOSIS was enriched for mitochondrial apoptosis and oxidative-stress–associated genes (e.g., *Bax* and ROS-handling/mitochondrial components, such as *Vdac2*, Gpx*1/3*, and *Sod2*)^58–61^, whereas progenitors preferentially upregulated stress-adaptive and pro-survival factors (e.g., *Mcl1*, *Ier3*, and *Gadd45b*) ^62–64^ (**Supplementary Fig. 3e**). EMT-related genes further suggested distinct remodeling modes, with progenitors biased toward adhesion/basement-membrane components (e.g., *Col4a1*/*Col4a2*, *Lama2*, and *Sparc*)^65–67^ and atretic cells showing increased collagen-processing and ECM-remodeling programs (e.g., *Plod2*, *Serpinh1*, and *Htra1*)^68–70^ (**Supplementary Fig. 3f**).

### Spatial deconvolution of GC subtypes at single-cell resolution by Stereo-seq

Although scRNA-seq enables high-resolution characterization of GCs transcriptional states and subtypes, it lacks spatial coordinates and thus cannot resolve where each subtype resides within ovarian tissue, how they are organized around follicular architecture, or how cellular composition and gene-expression patterns vary across key follicular regions^71^.

To connect scRNA-defined GC subtypes to spatial context, we analyzed a publicly available Stereo-seq FF V1.3 dataset of C57 mouse ovaries at a comparable age (6–8 weeks) with single-cell–resolution cell bin segmentation. We applied cell2location for spatial mapping at the cell bin level and assigned each cell bin to the most likely GC subtype based on posterior estimates. We interpreted the mapped patterns jointly with H&E morphology (**Fig. 4f, g and Supplementary Fig. 4**)^72^. Overall, the spatial mapping results were consistent with follicle distribution inferred from histology: all seven GC subtypes identified from scRNA-seq could be reliably mapped onto follicular structures and exhibited clear regional specificity and spatial layering (**Fig. 4f, g**). For visualization and validation, we selected representative follicular regions on the H&E section (α–θ) and cross-validated the mapped subtypes using the spatial distribution of canonical markers (**Fig. 4h**).

In region α, cell2location spatial mapping indicated a predominance of pGCs. The corresponding H&E morphology suggested a follicle surrounded by a single GC layer, consistent with a primary follicle, and the spatial enrichment of *Pcsk6* and *Wt1* with weak *Vcan* signal matched the *Wt1*^+^*Vcan*^−^ signature of progenitor cells. In region β, follicles were primarily composed of Preantral 1, Preantral 2, and Mitotic 1 cells and displayed a layered organization, with Preantral 1/2 distributed around the follicle and Mitotic 1 positioned closer to the oocyte side; histology suggested a secondary follicle–like stage. Notably, the Mitotic 1 signal was spatially biased toward the oocyte-adjacent compartment, suggesting that proliferative activity may be relatively enriched on the inner side at this stage. The *Wt1*^+^*Vcan*^+^ spatial pattern in this region was concordant with the preantral transcriptional features defined in scRNA-seq, supporting the utility of these signatures for distinguishing progenitor versus preantral GCs.

Region γ corresponded to a typical antral follicle on H&E, and spatial mapping recapitulated this architecture: mural GCs localized predominantly to the outer follicular wall, whereas the inner cumulus compartment was biased toward Mitotic 2. Marker distributions further supported this arrangement, with *Inhba* enriched in the outer mural layer and *Inhbb* detected in both compartments but biased toward the inner side. In addition, *Top2a* and *Kctd14* signals were observed in the inner compartment, suggesting a population that combines proliferation-associated features with cumulus-related markers, consistent with the scRNA-seq–inferred pseudotime trajectory of GC maturation. Finally, region θ corresponded to an atresia-associated follicular structure. Spatially, *Top2a* and *Inhbb* signals were still detectable near the central cumulus region but were overall weaker, whereas atresia-associated genes, such as *Grb14* and *Prss23*, were stronger in the outer region. Concordantly, spatial mapping suggested a more cumulus-like identity centrally and an Atretic GC-dominated outer compartment, consistent with an “outside-to-inside” progression of atresia-associated states. We also observed Atretic GC distributions around follicles of varying sizes, indicating that entry into an atresia-associated transcriptional state is not restricted to a single developmental stage.

Together, single-cell–resolution spatial mapping provides independent spatial support for the marker-defined GC subtypes and reveals their layered organization within follicles. These results establish a spatial framework for subsequent analyses of stage- and region-dependent oocyte–GC interactions.

### Identification of stage-resolved oocyte–GC communication

Guided by the cell2location-based spatial mapping, we constructed stage-matched oocyte–GC pairing groups by virtually pairing oocyte stages with their most likely surrounding GC subtypes, and then built CellChat objects to infer intra-follicle communication (**Fig. 5a**)^73^. Specifically, EGO oocytes were paired with pGCs; GO1 oocytes were paired with Preantral 1/2 and Mitotic 1; GO2 oocytes were paired predominantly with Mitotic 1/2; and the more advanced GO3 and FGO oocytes were paired with Mitotic 2 and Antral Mural GCs, consistent with their antral-like positioning in the mapped follicular context.

**Fig. 5.**
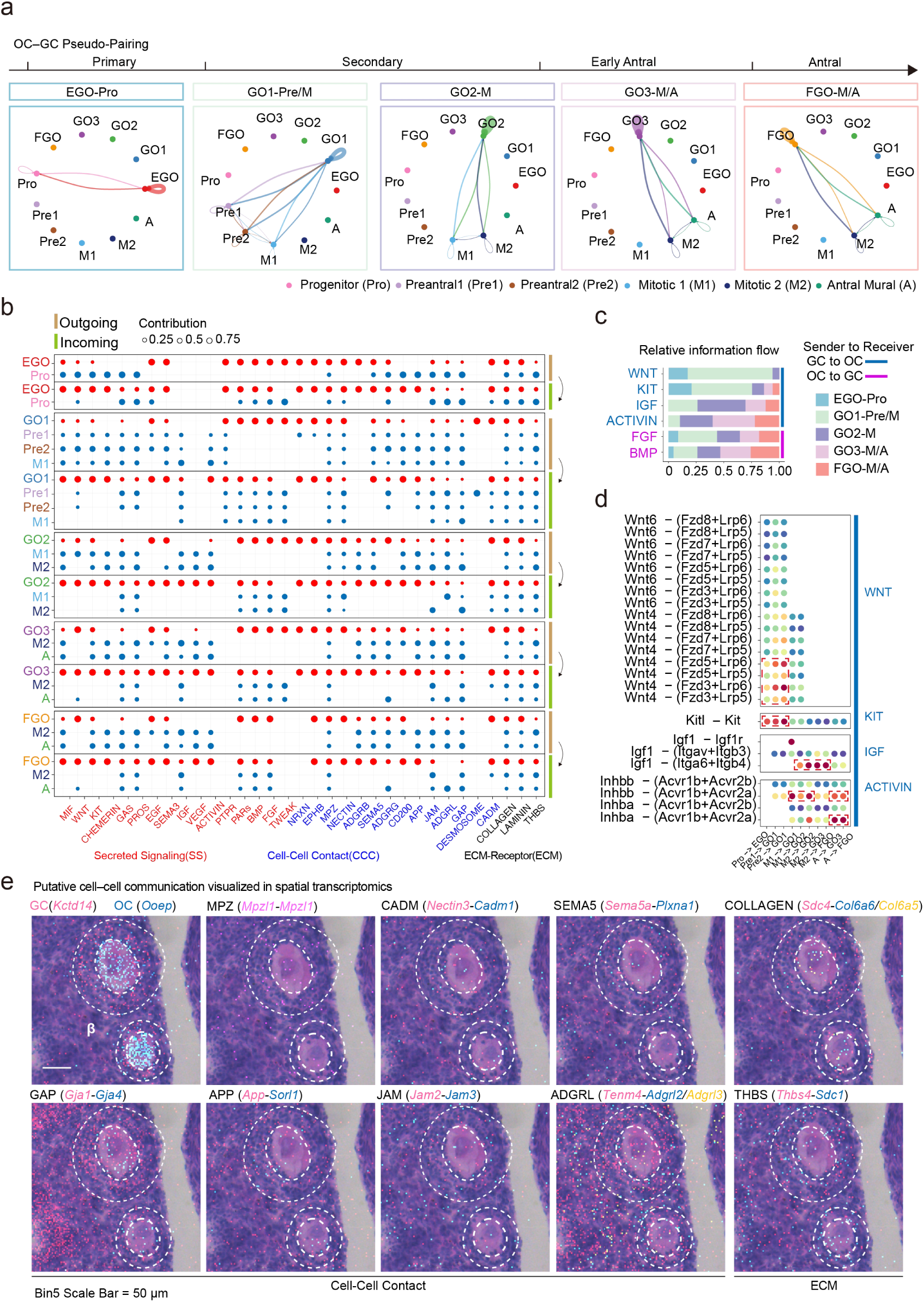
Putative intra-follicle oocyte–GC signaling inferred across oocyte growth stages. **a**, Stage-matched oocyte–GC pairing groups across the developmental progression (top); curves indicate predominant inferred pairings between oocyte and GC states. **b**, Outgoing (sender) and incoming (receiver) contributions of inferred oocyte–GC signaling pathways, grouped into secreted signaling (SS), cell–cell contact (CCC), and ECM–receptor (ECM); dot size indicates pathway contribution strength. **c**, Relative information flow of selected pathways across stages with inferred directionality (GC→oocyte or oocyte→GC). **d**, Representative ligand–receptor pairs for WNT, KIT, IGF, and ACTIVIN signaling across oocyte–GC pairs; dot size and color denote communication strength. **e**, Spatial transcriptomics (region β) showing representative expression patterns of selected ligand–receptor genes from CCC and ECM pathways in follicles (dashed outlines indicate follicle boundaries). Scale bar, 50 μm.

We next estimated communication probability within each stage-matched group using computeCommunProb, applying the triMean method to obtain a robust group-level expression estimate and thereby prioritize fewer, higher-confidence interactions. We then focused on oocyte–GC cross-talk and, after filtering, retained 33 oocyte–GC signaling families spanning three major modalities—secreted signaling (SS), cell–cell contact (CCC), and ECM–receptor (ECM). We summarized their inferred outgoing and incoming contributions across stages (**Fig. 5b** and ligand–receptor pairs are provided in **Supplementary Fig. 5a**).

Within SS, the oocyte–GC interactome comprised distinct functional classes, including canonical follicle growth/differentiation cues (KIT, BMP, EGF, and ACTIVIN), metabolism-linked signaling (IGF), vascular/structural programs (VEGF), remodeling/protease-associated pathways (PARs and TWEAK), and stress/immune-like homeostatic signals (MIF, CHEMERIN, and GAS family signals)^74–77^. Interestingly, we found many SS pathways were highly directional, with ligands largely confined to one compartment and receptors enriched in the other (GC→oocyte or oocyte→GC). To quantify how these directional pathways are deployed across development, we compared their relative information flow across stages and observed a clear stage bias (Fig. 5c). Early stage-matched oocyte–GC pairing groups were dominated by GC→oocyte WNT and KIT signaling: WNT flow rose sharply at GO1, with GC-derived ligands dominated by *Wnt4* (with minor *Wnt6*) engaging oocyte Fzd–Lrp receptor pairs, and the Kitl–Kit axis showed a similarly early-biased profile (Fig. 5c, d). Together, these early-peaking pathways suggest close involvement in the primary-to-secondary follicle transition. In contrast, later stage-matched oocyte–GC pairing groups exhibited increased IGF, ACTIVIN, FGF, and BMP signaling, largely accompanied by progressive upregulation of granulosa-derived ligands such as *Igf1* and *Inhba*/*Inhbb* from mitotic intermediates toward the antral stage (**Fig. 5c, d**). Notably, *Inhba*/*Inhbb* (TGFβ superfamily) signal via oocyte Acvr receptors and SMAD2/3/4, temporally aligning with the rise of Smad3 regulon activity from GO2 and its maintenance through GO3–FGO, supporting *Inhba*/*Inhbb* as candidate upstream cues for Smad3-associated programs during late oocyte growth (**Fig. 3h and Fig. 5d**). Collectively, these SS pathways span diverse biological functions and reveal a temporally dynamic, stage-dependent rewiring of oocyte–GC paracrine communication during follicle development.

### CCC and ECM–receptor interactions in follicles

Beyond secreted cues, our CellChat analysis highlighted a set of CCC programs operating between oocytes and GCs. As expected, we found the gap-junction axis as a prominent and well-established communication route in follicles^78^. In addition, we detected several junctional-architecture related families at the transcriptomic level, including NECTIN, JAM, DESMOSOME, and MPZ/MPZL^79–81^, suggesting that, alongside gap junctions, oocyte–GC interfaces may engage a broader repertoire of adhesive/junctional linkages that are less well characterized in the follicular context but could contribute to cell–cell coupling. We also identified contact-dependent boundary/guidance cues (e.g., EPHB and SEMA5)^82,83^ and a group of synapse-like adhesion/receptor modules frequently described in neuronal or adhesion contexts (e.g., NRXN, ADGRL/ADGRG/ADGRB, and APP)^84–86^, expanding the candidate space of direct oocyte–GC coupling mechanisms inferred from scRNA-seq.

To obtain spatial support for these putative CCC interactions, we visualized spatial distributions of the paired genes for representative pairs in region β of the Stereo-seq dataset, where both oocytes and surrounding GCs showed robust transcript capture (**Supplementary Fig. 5c**). Using the spatial patterns of *Kctd14* (GC-enriched) and *Ooep* (oocyte-enriched), together with H&E morphology, we delineated the oocyte and its surrounding GCs within follicles. As a positive reference, the inferred GAP interaction showed the expected compartmentalization: *Gja1* signal was predominantly detected in surrounding GCs, whereas *Gja4* was enriched toward the oocyte compartment, consistent with the known asymmetric connexin usage at the oocyte–GC interface. In the same follicles, multiple additional CCC examples—MPZ (*Mpzl1*–*Mpzl1*), CADM (*Nectin3*–*Cadm1*), SEMA5 (*Sema5a*–*Plxna1*), APP (*App*–*Sorl1*), JAM (*Jam2*–*Jam3*), and ADGRL (*Tenm4*–*Adgrl2*/*Adgrl3*)—displayed spatial distributions consistent with the inferred sender–receiver compartmentalization, supporting the in-situ compartmentalization expected from the inferred directionality (**Fig. 5e**).

In addition to contact-mediated programs, we also observed prominent ECM–receptor interactions at the oocyte–GC interface. In the ovary, the follicular basal lamina is classically enriched for basement-membrane collagens (e.g., COL4) and laminins, providing structural support around the granulosa compartment. In our data, however, oocytes showed notable expression of non–basement-membrane collagen transcripts, particularly COL6 family members (e.g., *Col6a5*/*Col6a6*) and *Col9a3*, which were paired with syndecan and integrin receptors (e.g., *Sdc4* with *Itga9*/*Itgb1*) (**Fig. 5e**)^87^. This pattern suggests that oocyte-derived ECM components may contribute to integrin/syndecan-dependent adhesion and signaling within follicles, potentially linking local matrix cues to oocyte–GC interface organization. We also detected strong GC-enriched *Thbs4* together with oocyte-enriched *Sdc1*, consistent with a THBS–syndecan axis that can modulate adhesive microenvironments and receptor-associated signaling at the oocyte–GC boundary (**Fig. 5e**)^88^.

Together, these CCC and ECM–receptor signals expand the candidate repertoire of direct oocyte–GC coupling beyond canonical gap junctions, spanning multiple adhesion, guidance, and matrix-sensing programs. In spatial transcriptomics, compartment-enriched and interface-adjacent expression patterns for representative paired genes are consistent with the inferred directionality of these interactions, providing spatial support for putative contact- and matrix-mediated coupling in situ (**Fig. 5e**).

## Discussion

In our previous work, we established that Stereo-cell can profile transcriptomes from large oocytes at scale. Here, we refined oocyte pretreatment, which enables more reliable joint interpretation of morphology and transcriptional state. By coupling imaging with transcript capture in the same cell, this framework helps anchor transcriptional states to biologically interpretable morphological status, rather than relying on transcriptome alone as a proxy for maturation.

A key bottleneck in the field is staging the “growing” continuum. Unlike primordial oogenesis (small cells) or FGO–MII comparisons (often synchronized by hormones), intermediate growth stages span large size changes and continuous heterogeneity and are commonly staged with unavoidable manual bias. By combining unsupervised transcriptome clustering with cell morphology, we provide a high-throughput framework to assign biologically interpretable stages across the maturation process with minimal intervention. This dual-modality staging offers an internal cross-check that conventional scRNA-seq alone cannot easily provide and increases temporal resolution across successive windows of maturation. Conceptually, this allows the growing-to-mature trajectory to be partitioned into higher-resolution sequential windows that can be compared across datasets and linked to specific developmental events. Using this framework, we identified an early window (EGO) with peak nuclear-encoded mitochondrial gene expression and oxidative phosphorylation programs, suggesting an early surge in mitochondrial energy production programs as growth initiates. A plausible interpretation is that primary-follicle–like oocytes transiently elevate mitochondrial biogenesis/respiratory capacity to meet the demands of rapid cytoplasmic growth and to prepare for later competence acquisition. Later stages showed a shift toward post-transcriptional regulation and cellular remodeling, consistent with acquisition of meiotic competence.

Because oocyte growth depends on the follicular microenvironment, we also profiled the matched ovarian somatic fraction and capturing nearly all major somatic cell types. Focusing on GCs, we reconstructed subtype relationships and identified a progenitor-like subset within mouse preantral GCs that follows a maturation trend similar to that described in human ovaries. Notably, although *Wt1* has been proposed as a marker of theca progenitors in mouse, we found that *Wt1* expression was not restricted to theca progenitors and was also high in progenitor and preantral GC subsets. Single-cell–resolution spatial strategies that integrate accurate cell segmentation with neighborhood-based microenvironment modeling have proven powerful for resolving local niches^89,90^, yet remain rarely applied at comparable resolution in the ovary. To place these subtypes in tissue context, we leveraged a Stereo-seq dataset with single-cell–resolution segmentation. Interestingly, mitotic-like GC states reported across studies preferentially localized to the peri-oocyte region in follicles, resembling a cumulus-like compartment, and further separated into two maturation-graded subtypes (Mitotic 1→Mitotic 2), consistent with a transitional continuum toward the antral state. This spatial bias suggests that proliferative GC programs may be coupled to an oocyte-proximal niche during the secondary-to-antral transition, providing a concrete tissue context for interpreting “mitotic” GC annotations across studies.

Finally, leveraging spatial mapping, we assembled stage-matched oocyte-GC pairing and summarized oocyte–GC communication across development, revealing stage-biased activation of canonical pathways (e.g., early WNT/KIT versus later ACTIVIN/IGF) together with additional contact- and ECM-linked candidates. These inferred programs offer a working model in which distinct paracrine, adhesive, and matrix-sensing signals are deployed in a temporally ordered manner as follicles progress through growth and antral formation. Although this virtual pairing does not track true follicle-matched oocyte–GC pairs, it provides a structured reference set of candidate interactions for subsequent spatial and functional follow-up. Future follicle-resolved profiling and perturbation experiments will be important to establish causal relationships and to validate directionality at the protein and signaling-activity levels. Overall, our work refines an oocyte-oriented dual-modality platform and provides an integrated atlas of oocyte and somatic programs across maturation, offering a resource and framework for studying follicle development and related disorders.

## Materials and Methods

### Sample collection

All animal procedures were performed in compliance with the guidelines for the care and use of laboratory animals and were approved by the Laboratory Animal Experimental Ethical Inspection of Dr. Can Biotechnology (Zhejiang) Co., Ltd. (approval number: 2024DRK0019). Female C57BL/6J mice at postnatal day (P)14, P21, and P49 were purchased from QIZHEN Laboratory Animal Technology Co., Ltd. (Hangzhou, China).

#### Collection of growing oocytes and ovarian somatic cells

Ovarian single-cell suspensions were prepared based on a published protocol with minor modifications^16^. Briefly, ovaries were dissected and minced into ∼0.5 mm³ cubes. The tissue was digested in 1 mg/mL type IV collagenase (BBI Life Sciences Corporation) at 37 °C with shaking at 1000 rpm in a Thermomixer for 20 min. Cells were then pelleted by centrifugation at 300 × g for 3 min at room temperature, and the supernatant was removed. The pellet was resuspended in TrypLE Express (Thermo Fisher Scientific) and incubated at 37 °C with shaking at 1000 rpm for 7 min. Digestion was quenched with M2 medium (Sigma-Aldrich), and any undigested residual tissue was manually removed, yielding an ovarian single-cell suspension.

Intact, visually identifiable denuded oocytes without adherent GCs were collected from the suspension using mouth pipetting and transferred into M2 medium droplets under mineral oil on a 37°C warming stage for immediate downstream pretreatment. After oocyte collection, the remaining single-cell suspension was passed through a 40 μm cell strainer and processed promptly for droplet-based scRNA-seq library preparation.

#### Collection of hormonally primed MII oocytes

For MII oocyte collection, the protocol was followed as described. P21 females were injected with 5 IU PMSG (Sigma-Aldrich), followed 46 h later by 5 IU hCG (Sigma-Aldrich). Cumulus–oocyte complexes (COCs) were harvested from the ampulla of the oviduct 15 h after hCG injection. Cumulus cells were removed by incubation in hyaluronidase droplets (Sigma-Aldrich), and oocytes were washed thoroughly in M2 medium. Clean denuded MII oocytes were transferred to fresh M2 medium droplets under mineral oil on a 37°C warming stage and processed immediately for downstream pretreatment.

### Stereo-cell oocyte pretreatments

Oocytes in M2 medium droplets were transferred into Tyrode’s solution (Sigma-Aldrich) droplets to remove zona pellucida. During zona pellucida removal, the dish was gently swirled continuously, while oocytes were monitored in real time under a microscope. Immediately after complete zona dissolution, oocytes were transferred into fresh M2 droplets for thorough washing.

Zona-free oocytes were then moved into dishes containing pre-cooled 50% methanol (v/v in PBS) to initiate stepwise methanol fixation. Every 3 min, an equal volume of pre-cooled 100% methanol was added with gentle mixing to increase the methanol concentration; this step was repeated three times. Subsequently, most of the solution was carefully removed (avoiding oocyte loss), replaced with 100% methanol, and oocytes were fixed for an additional 10 min.

### Stereo-cell library preparation, sequencing

Stereo-cell chip preparation and library construction were carried out largely according to our previously reported protocol^14^. In brief, Stereo-seq chips were PLL-precoated (Sigma-Aldrich) before sample loading. Pre-treated oocytes were resuspended in pre-chilled 100% methanol and loaded onto the PLL-precoated chip in multiple rounds (typically 20–30 oocytes in 10 μL methanol per round). Each round was followed by brief air drying to allow methanol evaporation and stable adhesion of oocytes to the chip surface, after which the next aliquot was applied. For a single S1 chip (1 cm × 1 cm), this procedure was usually repeated 5–7 times. After the final loading, the chip was gently air-dried to further stabilize oocyte–chip contact.

After loading, subsequent steps followed the standard Stereo-cell workflow, including DAPI staining (Beyotime Biotechnology) and imaging for morphology and spatial registration, permeabilization and probe hybridization, on-chip reverse transcription to generate cDNA, and enzymatic cDNA release followed by bead-based cleanup and PCR amplification for downstream library preparation. DNA nanoballs (DNBs) were produced and sequenced on the MGI DNBSEQ-Tx system using the same read configuration as in our previous method (50 bp for read 1 and 100 bp for read 2).

### Stereo-cell raw data preprocessing

Raw sequencing reads were processed to obtain CID-resolved, UMI-collapsed gene counts. Specifically, the CID sequence in read 1 was matched to a predefined coordinate barcode whitelist, allowing up to one nucleotide mismatch. Reads were retained only when both the CID was valid and the UMI passed quality filters (UMIs containing no more than two bases with Phred ≤10 and no ambiguous “N”). Qualified reads were converted to a FASTQ+ representation in which the CID and UMI information was incorporated into the read header. The resulting reads were aligned to the mouse reference genome (mm10) with STAR^91^; only uniquely aligned reads were used for annotation and counting, and alignments with a mapping score <10 were removed. Finally, UMIs were deduplicated per gene × CID, and outputs were summarized as a spatially resolved gene expression matrix (sGEM) file comprising gene ID, x coordinate, y coordinate, total UMI count, and intronic UMI count.

### Stereo-cell staining image registration

Registration between the DAPI staining image and the UMI signal was performed using TrakEM2 in ImageJ (v1.53f51). Briefly, the spatial gene expression matrix was first converted into a grayscale image representing the UMI distribution. This UMI image, together with the staining image, was imported into a new TrakEM2 project. After unlocking the image layers, the UMI image was assigned to the red channel and the staining image was assigned to the green channel. The green channel was then manually adjusted (translation and scaling) until it visually overlapped with the red channel. The aligned green channel, corresponding to the registered staining image, was finally exported and saved as a new registered flat image for downstream analysis^92,93^.

### Stereo-cell oocyte segmentation and generation of cell-resolved expression matrices

Registered DAPI–UMI images were imported into QuPath (v0.4.3) for manual oocyte segmentation^94^. For each field, the UMI grayscale channel and the DAPI channel were compared between channels to ensure that each selected UMI pattern corresponded to a single DAPI-stained oocyte. Using QuPath annotation tools, only well-isolated oocytes with one-to-one UMI–DAPI correspondence were outlined on the UMI channel following the UMI signal boundary (one ROI per oocyte). Regions showing overlapping UMI signals from multiple cells or UMI-positive areas without a matching DAPI-stained oocyte were excluded. All ROI objects were then exported as a GeoJSON file.

Oocyte diameters were measured in QuPath using the DAPI channel. DAPI staining produced a detectable intracellular background, and image contrast was adjusted to make the oocyte boundary visible. The line tool was then used to draw a diameter line within each oocyte ROI, and the corresponding length measurement was recorded for downstream analyses.

To extract cell-specific transcript counts, the GeoJSON ROIs were converted into sets of coordinates within each ROI, and the original GEM file was filtered accordingly by assigning molecules to ROIs to generate a cell-labeled GEM. Finally, a sparse gene-by-cell UMI count matrix was constructed.

### DNBelab C4 library construction, sequencing and data processing

Ovarian single-cell suspensions were converted into droplet-based scRNA-seq libraries using the DNBelab C4 workflow (MGI, 940-001818-00). Briefly, 15,000–20,000 viable single cells were loaded onto the DNBelab C-TaiM 4 system (MGI, 900-000637-00) for cell partitioning, barcoding, and reverse transcription, following the manufacturer’s instructions. Barcoded cDNA was subsequently recovered and fragmented to generate sequencing-ready libraries. Libraries were sequenced on a DNBSEQ-Tx platform with a paired-end configuration (50 bp for read 1 and 100 bp for read 2). Sequencing data were processed with DNBC4tools (https://pypi.org/project/DNBC4tools/) to obtain the gene-by-cell expression matrix. All read alignment and gene annotation were performed using the mm10 mouse reference genome.

### Single-cell data quality control and analysis

#### Stereo-cell

Stereo-cell gene-by-cell UMI count matrices were used to create Seurat objects for downstream analyses^95^. The following describes the Seurat workflow: (1) SCTransform was applied to each library for normalization and identification of highly variable genes (HVGs). Common HVGs across libraries were then selected using SelectIntegrationFeatures. (2) All Stereo-cell libraries were merged, and SCTransform was re-run on the merged object for normalization and scaling, with ‘residual.features’ set to the common HVGs. (3) Principal component analysis was performed using RunPCA. (4) Batch effects were corrected with RunHarmony^96^. (5) Cell–cell neighborhoods and community detection were carried out using FindNeighbors and FindClusters. (6) Two-dimensional visualization was generated with RunUMAP.

#### DNBelab C4

The DNBelab C4 gene-by-cell expression matrices were used to create Seurat objects for downstream analyses. The overall analysis pipeline was similar to that used for Stereo-cell, with additional preprocessing steps. Briefly, ambient RNA contamination was removed using SoupX^97^, and potential doublets were identified and excluded using scDblFinder^98^. NormalizeData, FindVariableFeatures, and ScaleData (instead of SCTransform) were then used for normalization, HVG selection, and scaling. For granulosa-cell subclustering, cell-cycle effects were regressed out during scaling (using S and G2/M scores) to reduce cell-cycle–driven variation. Dimensionality reduction, batch correction, clustering, and UMAP visualization were performed as described above.

### Trajectory inference

Oocyte developmental trajectories were inferred using scTour (v1.0.0) and Monocle3 (v1.0.0). For scTour, a feature set of stage-specific marker genes was defined and identified using Seurat FindAllMarkers (default parameters). The scTour model was trained using mean squared error (MSE) loss, and a vector field was constructed in the scTour-derived latent space to characterize transitions between cell states, thereby deriving a continuous ordering of cells along the developmental trajectory.

Pseudotime provides a quantitative coordinate to order single cells along a continuous biological process and enables downstream analyses of gene expression changes along progression. For Monocle3, the Seurat object was converted to a cell_data_set (cds) and augmented with Seurat-derived UMAP coordinates, cluster labels, and stage annotations. All cells were assigned to a single partition (partition = 1). A principal graph was learned in the fixed UMAP space using learn_graph, and pseudotime values were computed using order_cells. Cells were then evenly binned into 13 pseudotime bins based on their pseudotime values. Temporal-sensitive genes were identified using graph_test with the criteria q_value < 0.05 and morans_I > 0.2. These genes were further grouped into five modules using find_gene_modules (resolution = 0.005).

### Developmental potential analysis

Developmental potential was assessed using CytoTRACE2 (v1.0.0). The mouse model was applied to calculate a developmental potential score for each oocyte.

### pySCENIC

We employed pySCENIC (v0.12.1) to infer gene regulatory networks (GRNs) in oocytes and assess transcription factor (TF) activities at the single-cell level. First, GRNBoost2 was used to infer co-expression modules between transcription factors and candidate target genes, using as input a loom file containing oocyte expression data restricted to Monocle3 pseudotime bins 1–9 (excluding bins 10–13 enriched for MII oocytes, which lack de novo transcription). Next, the cisTarget pipeline was utilized to prune the initial co-expression network by identifying motif-supported regulatory interactions, resulting in the final regulon network. Finally, regulon activities were quantified across individual cells using AUCell scoring, yielding an AUC score for each regulon. These analyses required a set of predefined reference resources, including a transcription factor list, motif rankings database, and motif-to-TF annotation file. The motif rankings and motif to TF annotations used in this study were taken from the v10 (mc_v10_clust) gene-based 10 kb cis regulatory database for the mm10 mouse genome assembly. All reference files used in this study were obtained from the publicly available resources provided by the Aerts lab (https://resources.aertslab.org/cistarget/databases/).

### Enrichment analysis

#### GO analysis

Gene ontology (GO) enrichment was performed in R using clusterProfiler with the mouse annotation package org.Mm.eg.db^99^. Candidate gene sets (e.g., pseudotime-associated genes or marker genes identified by Seurat FindAllMarkers) were tested for Biological Process (BP) enrichment using enrichGO. Redundant GO terms were further reduced using the simplify function to obtain a non-redundant set of representative BP terms.

#### ssGSEA analysis

Single-sample gene set enrichment analysis (ssGSEA) was used to quantify pathway/signature activity across predefined groups (e.g., granulosa-cell subtypes or oocyte stages). The ssGSEA scores were computed with GSVA using group-level average expression matrices as input. Gene sets were derived from MSigDB Hallmark collections (for granulosa-cell functional activity) and from stage-specific oocyte signature gene sets defined in this study.

### Cell–cell communication analysis

Cell–cell communication was inferred using CellChat (v2.1.2). GC and oocyte Seurat objects were merged, and the normalized expression matrix and cell annotations (GC subtypes and oocyte stages) were extracted from the merged object. To enable stage-resolved analyses, cells were split into five datasets (EGO, GO1, GO2, GO3, and FGO), each comprising oocytes from the corresponding stage together with the GC subtypes present in the same dataset. For each dataset, a CellChat object was constructed from the normalized expression data and analyzed with the CellChatDB.mouse ligand–receptor database. Communication probabilities were inferred using the standard CellChat pipeline, and signaling pathways were summarized to generate pathway-level communication networks. For cross-stage comparisons, cell groups were harmonized with liftCellChat, stage-specific CellChat objects were integrated with mergeCellChat, and differential communication patterns were visualized using rankNet.

### Stereo-seq data analysis

#### Raw data preprocessing

Publicly available mouse ovary Stereo-seq Transcriptomics FF v1.3 demo data were obtained from the STOmics website (https://www.stomics.tech/col1347). Raw FASTQ files were processed using the Stereo-seq Analysis Workflow (SAW), including spatial barcode mapping, alignment of cDNA reads to the mm10 mouse reference genome using STAR, and generation of a gene-by-spot count matrix for downstream analyses. Single-cell–resolution “cell bin” matrices were generated from image-based single-cell segmentation of the ovary H&E image using deep-learning–based CellBin pipelines.

#### Data processing and spatial gene visualization

Downstream analyses were performed on the processed spatial expression matrices. Briefly, basic quality control was applied, followed by normalization, log-transformation, dimensionality reduction, neighborhood graph construction, and clustering. SAW-generated outputs were used for spatial visualization of gene expression in StereoMap V4.

#### Cell2location-based spatial mapping

To estimate cell-type distributions in space, we applied cell2location (v0.1.4) using our single-cell RNA-seq data as a reference. In the single-cell dataset, mitochondrial genes were removed and informative genes were selected. A reference regression model was trained to learn cell-type–specific expression signatures. After intersecting genes between the reference and spatial datasets, the spatial model was fit to infer cell-type abundances for each spatial location. Abundance estimates were stored in the AnnData object for visualization and downstream analyses. Because the demo ovary data was generated from 6–8-week-old female C57BL/6 mice, we used our age-matched P49 ovarian single-cell dataset as the reference for projection.

## Acknowledgements

This work was supported by the National Key Research and Development Program of China (2025YFC3409300 to Chuanyu L.; 2024YFA0919700 to Chang L.), Zhejiang Science and Technology Department (2025C01092 and 2024C03004), Hangzhou Science and Technology Department (2024SZD1B09; TD2023003; 2024SZD0128 to Chang L. and Chuanyu L.), Guangdong Basic and Applied Basic Research Foundation (2024B1515230003, 2026A1515012050 to Chuanyu L.), and the Shenzhen Key Laboratory of Single-Cell Omics (ZDSYS20190902093613831). We thank DCS Cloud (https://cloud.stomics.tech/) for providing the computational resources and software support necessary for this study. We thank all our team members and the Zhejiang Key Laboratory of Spatial Omics for their support. We also thank colleagues at BGI Research, including Liang Chen, Yue Yuan, Xiumei Lin, Qiuting Deng, Yingjie Luo, Yaling Huang, Xiaojing Xu, Yong Bai, Huan Hou, Hongyu Luo, Mengnan Cheng, Mingyue Wang, Zijie Ye, Xiaoqian Chen, and Yiting Lin, for support and helpful discussions across different stages of this work. We especially thank Yongjuan Sang (Zhejiang University), Shiya Cheng (Wuhan University), and Peng Yuan (Peking University) for insightful guidance on oocyte experimental techniques and project analysis.

## Author contributions

Chuanyu L., X.X. and Y.R. conceived the idea; Chuanyu L., Chang L., X.X. and Y.R. supervised the work; Chuanyu L., Chang L., S.Z. and Y.L. designed the experiments; S.Z., X.Y., R.L., J.Z. and J.L. performed the majority of the experiments; S.Z., H.W., and M.Y. performed data preprocessing and quality evaluation; S.Z. and Z.L. analyzed the data; Z.L. and H.X. performed image processing; Q.L., Z.Z., L.X. and X.Z. provided technical support; S.H. and Y.L. gave the relevant advice; S.Z. and Z.L. wrote the manuscript.

## Supplementary Figures

**Supplementary Figure 1.**
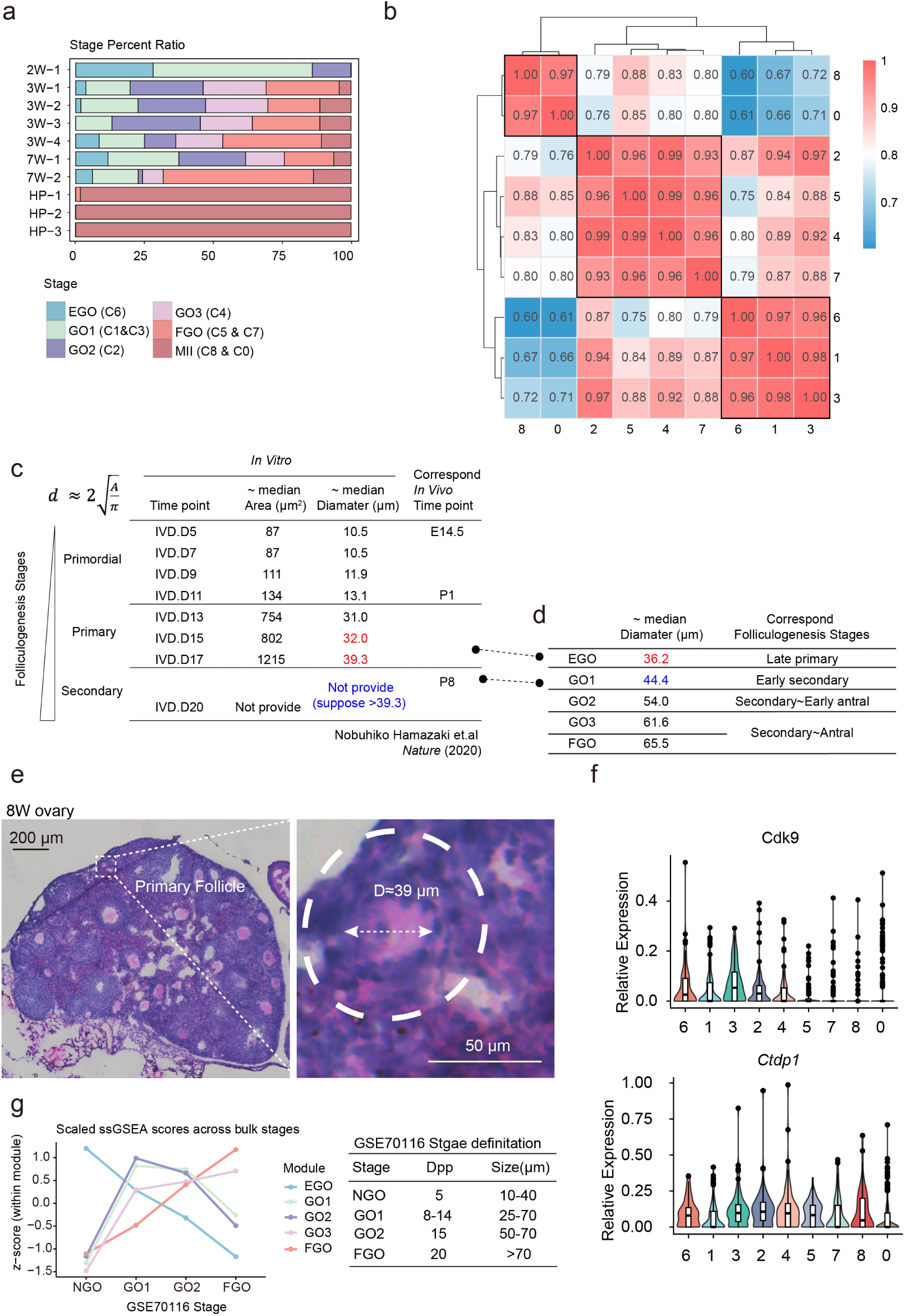
Supplementary analysis for oocyte stage annotation. **a**, Stage composition across individual Stereo-cell libraries shown as stacked bar plots (EGO, GO1, GO2, GO3, FGO, and MII). **b**, Pairwise Pearson correlation of transcriptomic profiles between clusters (C0–C8), shown as a correlation matrix. **c**, Summary of oocyte growth metrics from Hamazaki et al. (*Nature* 2020), including median oocyte area/diameter at each time point and the *in vitro* – *in vivo* correspondence reported by the authors. **d**, Mapping of stages defined in this study (EGO–FGO) to median oocyte diameters and corresponding folliculogenesis stages. **e**, Representative histology of an adult mouse ovary highlighting primary follicles and an example measurement of oocyte diameter. Scale bars, 200 μm (left) and 50 μm (right). **f**, Violin plots of *Cdk9* and *Ctdp1* expression across clusters. **g**, External validation using bulk RNA-seq dataset GSE70116: scaled ssGSEA scores of stage signature gene sets across bulk-defined oocyte growth stages (stage definitions shown at right).

**Supplementary Figure 2.**
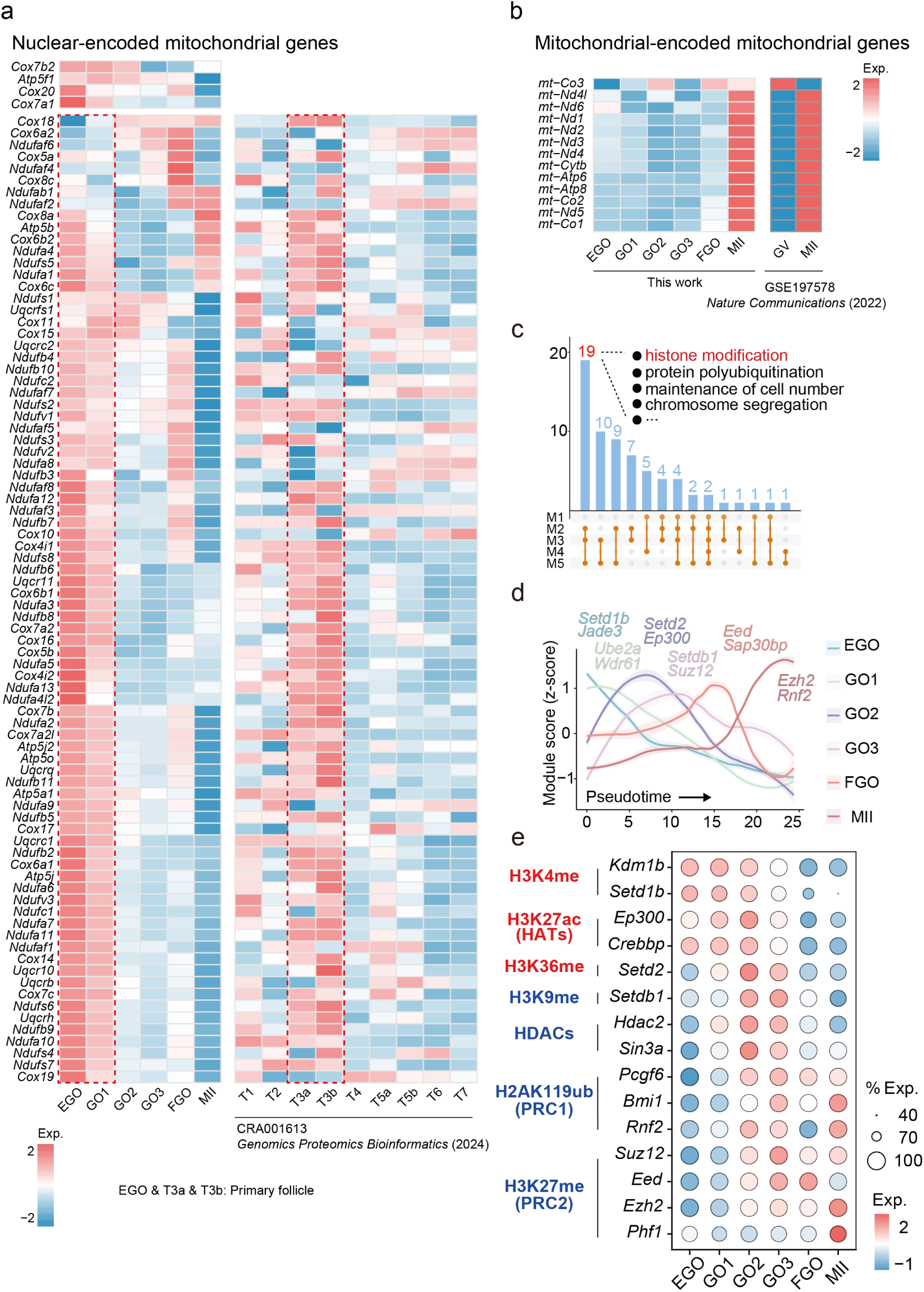
Additional evidence for stage-dependent metabolic and chromatin programs during oocyte growth and maturation. **a-b**, Heat maps show scaled expression (Exp.; row-wise z-scores) of nuclear-encoded mitochondrial genes (a) and mitochondrial genome–encoded genes (b) across stages in this study (EGO, GO1–GO3, FGO, and MII) and external datasets (CRA001613, T1–T7, GSE197578, GV, and MII). **c**, UpSet plot showing overlaps of significantly enriched GO terms among co-expression modules (M1–M5). Bars indicate the number of GO terms in each intersection. **d**, Module scores (z-scored) of histone modification–related gene sets plotted along pseudotime. Representative genes are labeled. **e**, Dot plot of representative histone modification–related regulators across stages (EGO, GO1, GO2, GO3, FGO, and MII), grouped by associated chromatin marks/complexes (left).

**Supplementary Figure 3.**
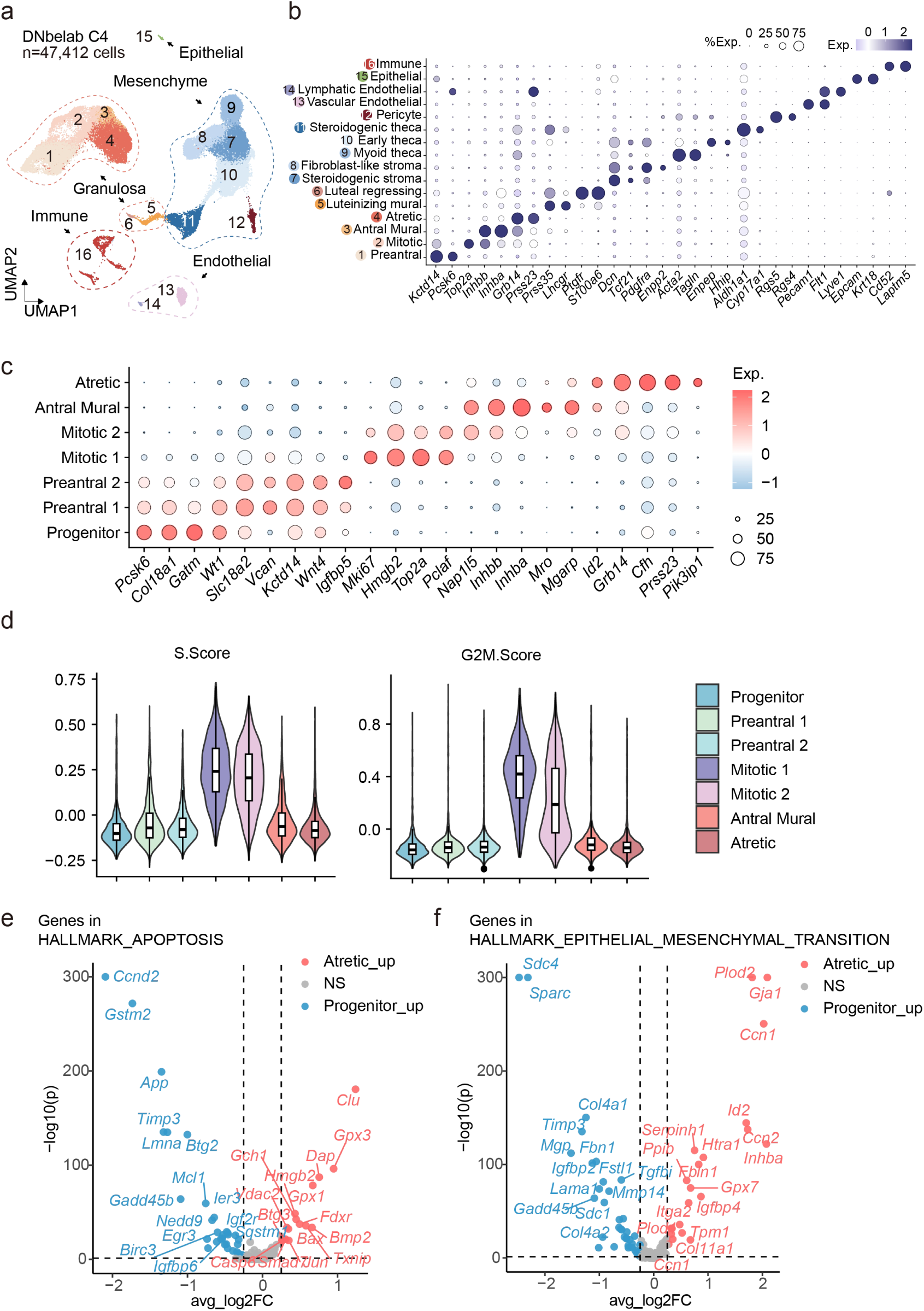
Ovarian cell-type annotation and granulosa subtype markers in scRNA-seq. **a**, UMAP of DNBelab C4 ovarian cells after quality control (n = 47,412), annotated into major cell types and subtypes. **b**, Dot plot of canonical marker genes used for major cell-type and subtype annotation. **c**, Dot plot of subtype-enriched markers across the seven granulosa subtypes. **d**, Violin plots of S-phase and G2/M cell-cycle scores across granulosa subtypes. **e**, Volcano plot of differential expression for genes in HALLMARK_APOPTOSIS between Progenitor and Atretic GCs; representative genes are labeled. **f**, Volcano plot of differential expression for genes in HALLMARK_EPITHELIAL_MESENCHYMAL_TRANSITION between Progenitor and Atretic GCs; representative genes are labeled.

**Supplementary Figure 4.**
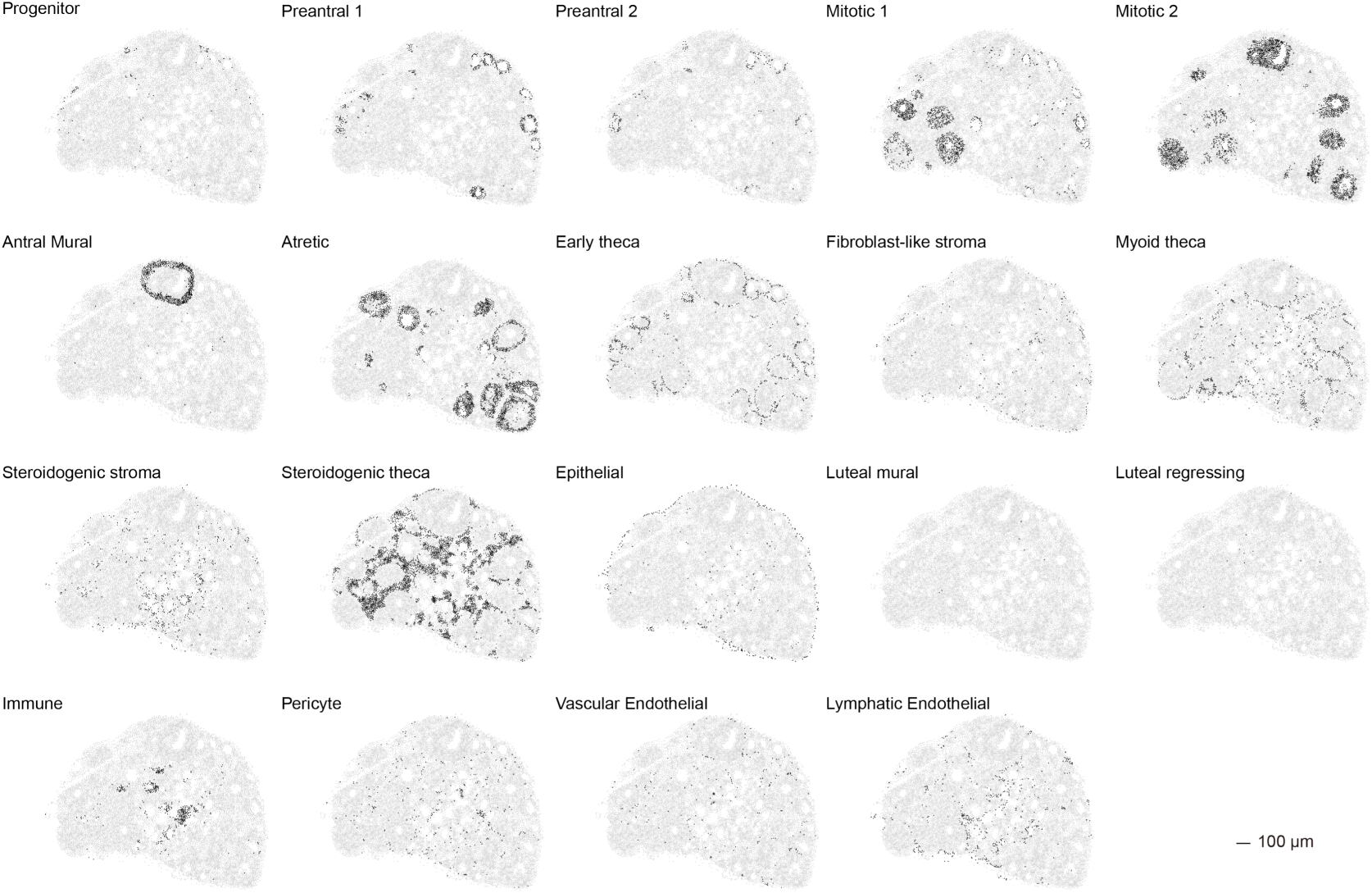
Cell2location mapping of ovarian cell subtypes in Stereo-seq FF ovary sections. Spatial maps showing cell2location-predicted localization of each annotated cell subtype on the Stereo-seq FF V1.3 ovary section (6–8 weeks old), displayed separately by subtype. Scale bar, 100 μm

**Supplementary Figure 5.**
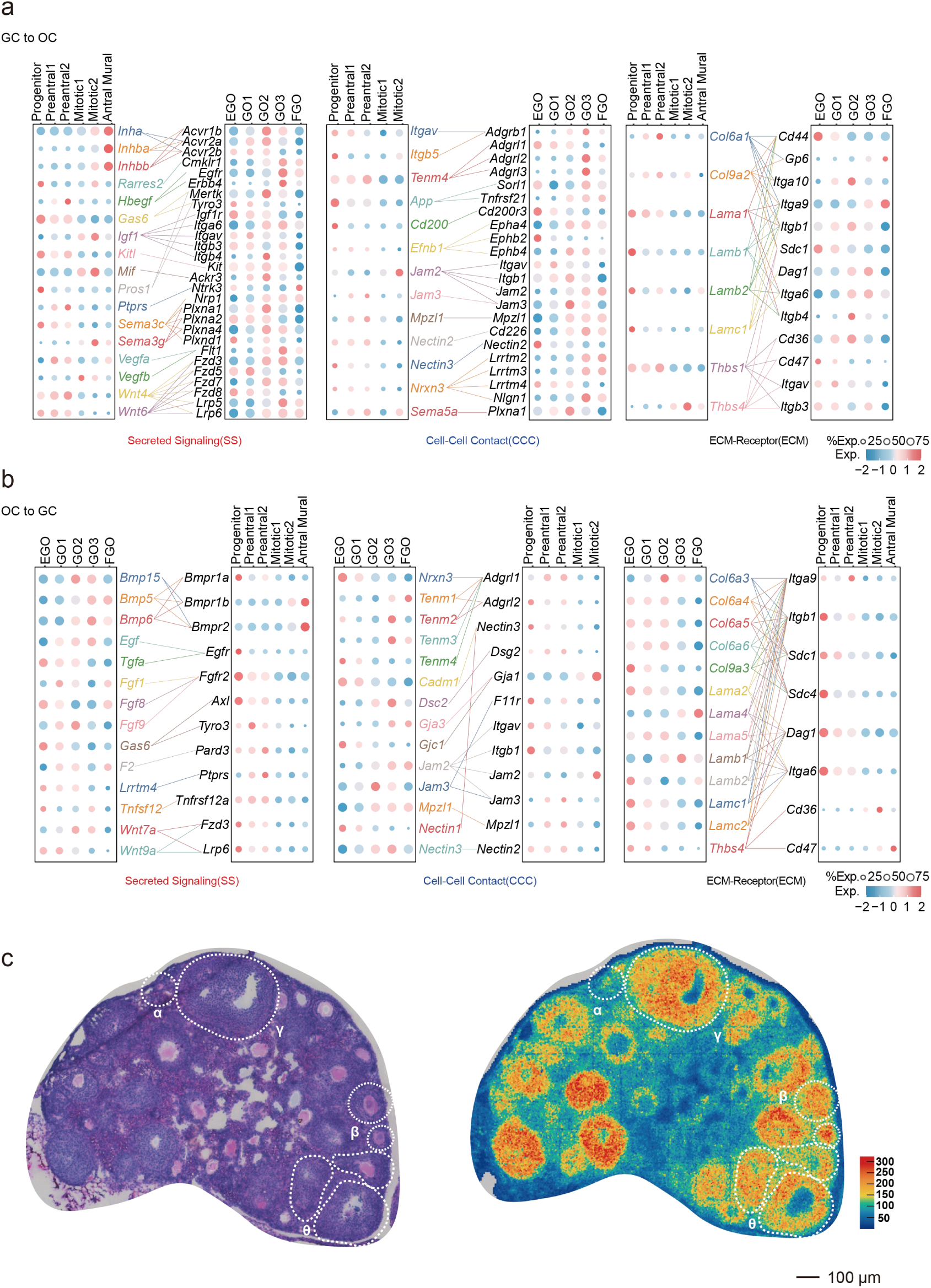
Expression of representative oocyte/GC ligand–receptor genes. **a-b**, Dot plots showing expression of representative ligand–receptor genes between GC subtypes and oocyte stages, grouped into secreted signaling (SS), cell–cell contact (CCC), and ECM–receptor (ECM), shown separately for GC→oocyte (a) and oocyte→GC (b); dot size indicates % expressing cells and color indicates scaled expression. **c**, Stereo-seq ovary section showing annotated follicle-enriched regions (left, H&E) and UMI density (right). Based on the UMI distribution, only two follicles in region β exhibited robust mRNA capture in the oocyte compartment and were therefore selected for spatial visualization of representative oocyte–GC ligand–receptor genes (dashed outlines indicate follicle boundaries). Scale bar, 100 μm.

